# Respiration-phased switching between sensory inputs and top-down inputs in the olfactory cortex

**DOI:** 10.1101/499749

**Authors:** Kimiya Narikiyo, Hiroyuki Manabe, Yoshihiro Yoshihara, Kensaku Mori

**Author notes:** Correspondence to: Kensaku Mori, Ph.D. Present address: Laboratory for Systems Molecular Ethology, RIKEN Center for Brain Science, 2-1 Hirosawa, Wako, Saitama 351-0198, Japan.

## Abstract

Olfactory perception depends on respiration phases: olfactory cortex processes external odor signals during inhalation whereas it is isolated from the external odor world during exhalation. Olfactory cortex pyramidal cells receive the sensory signals via bottom-up pathways terminating on superficial layer (SL) dendrites while they receive top-down inputs on deep layer (DL) dendrites. Here we asked whether olfactory cortex pyramidal cells spontaneously change the action modes of receiving olfactory sensory inputs and receiving top-down inputs in relation to respiration phases. Current source density analysis of local field potentials recorded in three different olfactory cortex areas of waking immobile rats revealed β- and γ-range fast oscillatory current sinks and a slow current sink in the SL during inhalation, whereas it showed β- and γ-range fast oscillatory current sinks and a slow current sink in the DL during exhalation. Sensory deprivation experiments showed that inhalation-phased olfactory sensory inputs drove the inhalation-phased fast oscillatory potentials in the SL but they drove neither the inhalation-phased slow current sink in the SL nor the exhalation-phased slow current sink in the DL. The results indicate that independent of inhalation-phased olfactory sensory inputs, olfactory cortex pyramidal cells spontaneously generate a slow depolarization in the SL dendrites during inhalation, which may selectively boost the concomitant olfactory sensory inputs to elicit spike outputs. In addition, the pyramidal cells spontaneously generate slow depolarization in the DL dendrites during exhalation, which may assist top-down inputs to elicit spike outputs. We thus hypothesize that the olfactory cortical areas coordinately perform inhalation/exhalation-phased switching of input biasing: inhalation phase is the time window for external odor signals that arrive in the SL dendrites, whereas exhalation phase is assigned to boost top-down signals to the DL dendrites that originate in higher brain centers.

## Introduction

Olfactory system is unique among sensory systems in that the sensory information processing in the higher olfactory centers is tightly coupled with respiration phases (Mori and Manabe, 2014). Olfactory sensory neurons in the nasal epithelium detect environmental odors and generate olfactory sensory signals only during inhalation phase of respiration, whereas they are physically isolated from the environmental odors during exhalation. On top of that, when mammals intend to explore the external odor world, they show the behavior of active inhalation to enhance odor inputs such as exploratory sniffing (Kepecs et al., 2007).

Olfactory sensory signals are transmitted along bottom-up olfactory pathways from the olfactory epithelium to the olfactory bulb and further to various areas of the olfactory cortex (Neville and Haberly 2004; Shepherd et al., 2004; Mori and Sakano, 2011; Mori and Manabe, 2014; Wilson and Sullivan, 2011). Because this transmission occurs in synchrony with inhalation (Manabe and Mori, 2013; Mori et al., 2013), we suppose that processing of olfactory sensory signals by the olfactory cortex does not occur continuously but takes place discretely in phase with inhalation.

The olfactory cortex consists of many areas including anterior olfactory nucleus (AON), ventral and dorsal tenia tecta, dorsal peduncular cortex, anterior piriform cortex (APC), olfactory tubercle, posterior piriform cortex (PPC), nucleus of the lateral olfactory tract, anterior cortical amygdaloid nucleus, posterolateral cortical amygdaloid nucleus, and lateral entorhinal cortex (Neville and Haberly, 2004). The AON, APC and PPC are major areas of the olfactory cortex and have a simple laminar structure with pyramidal cells as principal neurons. The pyramidal cells in these areas extend apical tuft dendrites to the superficial layers (SL dendrites in layers Ia and Ib) and receive olfactory sensory inputs in the SL either directly from axons of olfactory bulb neurons that terminate in layer Ia or indirectly from Ib-association fibers via a relay of pyramidal cells in the AON or APC (Fig. 8).

For the transformation of olfactory sensory inputs to appropriate odor-guided behaviors, the olfactory cortex requires not only bottom-up olfactory sensory signals from the external world but also top-down signals that are generated internally by higher level cognitive processes, such as attention, expectation, working memory and decision making (Gilbert and Sigman, 2007; Roelfsema and deLange, 2016; Shiotani et al., 2018). Pyramidal cells in the olfactory cortex extend proximal apical dendrites and basal dendrites in deep layers (DL dendrites in layers II and III of the APC and PPC, in layer II of the AON) and receive top-down inputs in the DL from the orbitofrontal cortex, ventral agranular insular cortex, medial prefrontal cortex and amygdala (Fig. 8) (Datiche and Cattarelli, 1996; Illig, 2005; Neville and Haberly, 2004). They also receive deep-association fiber inputs to the DL that originate from higher order areas of the olfactory cortex (Johnson et al., 2000; Chen et al., 2003; Neville and Haberly, 2004).

In the present study, we focused on the AON, APC, and PPC of awake rat with slow respirations and asked whether the pyramidal cells spontaneously change the action mode between actively receiving bottom-up olfactory sensory inputs and actively receiving top-down inputs in relation to inhalation and exhalation phases of respiration. We made multi-laminar electrophysiological recordings of local field potentials (LFPs) and single-unit spike activities in these areas with simultaneous monitoring of respiration. Current source density (CSD) analysis of LFPs in the AON, APC and PPC together with sensory deprivation experiments indicated that independent of inhalation-phased fast oscillatory olfactory sensory inputs, the pyramidal cells of these areas spontaneously generated slow and long depolarization in the SL dendrites during inhalation and slow and long depolarization in the DL dendrites during exhalation.

## Results

### Respiration-phased slow oscillatory activity in the olfactory cortex during wakefulness

To examine whether the timings of inhalation and exhalation are coupled with processing distinct synaptic inputs in the olfactory cortex, we recorded LFPs in the AON, APC or PPC using a linear microelectrode array together with the respiration pattern. In a well-acclimated head-fixed position, rats typically showed either awake state with slow respirations (0.5-3Hz) (Fig. 1 A-left, blue trace) or slow-wave sleep state with very stable slow respirations (0.5-2Hz) (Fig. 1 A-center, blue trace). These rats only occasionally displayed awake state with transient fast respirations (3-7 Hz) (Fig. 1 A-right, blue trace).

**Figure 1.**
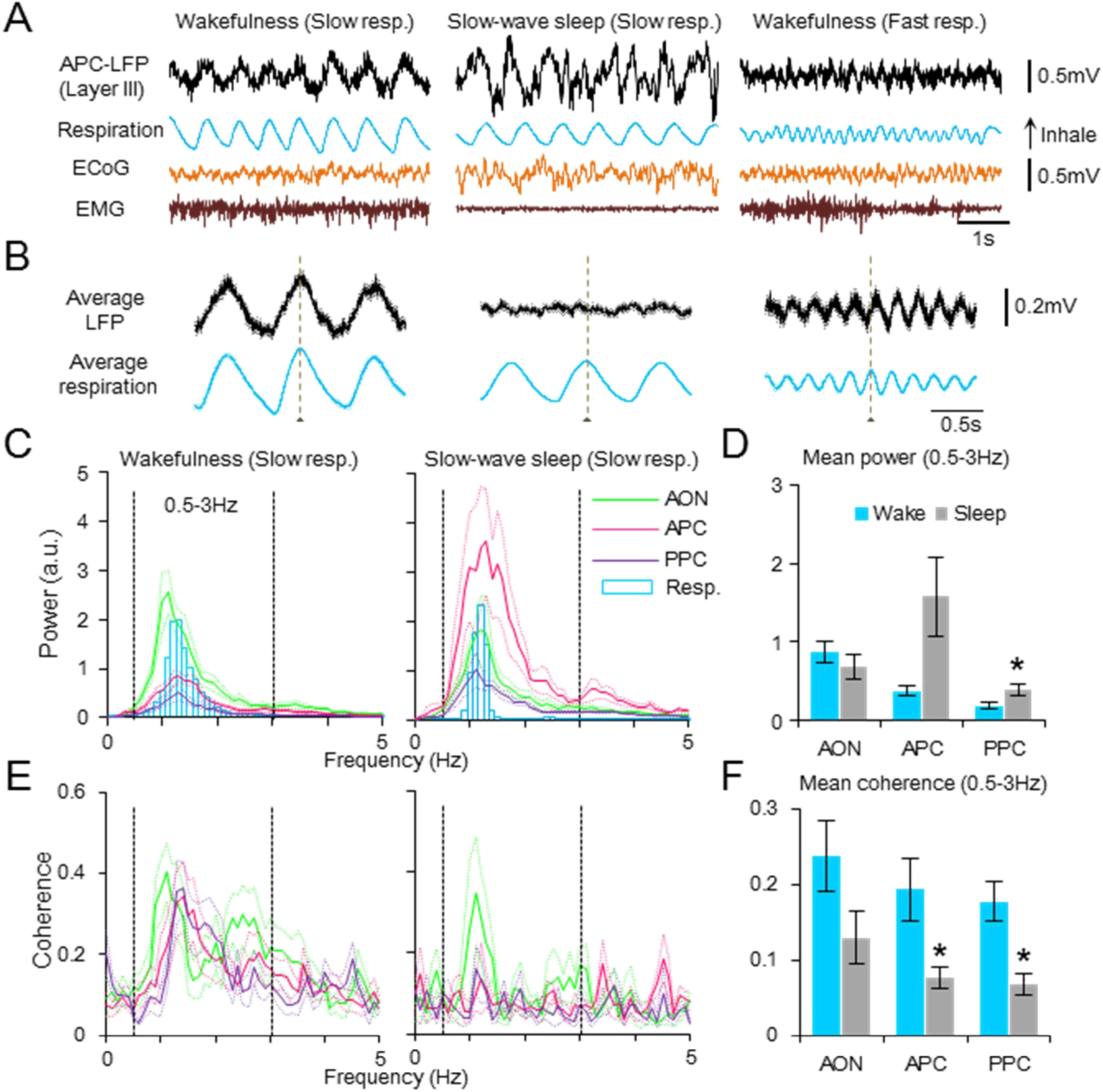
Respiration-phased slow oscillatory activity in the olfactory cortex. (A) A representative example of local field potential (LFP) recorded in the layer III of APC with simultaneously recorded respiration (blue trace), neocortex electrocorticogram (ECoG) and neck muscle electromyogram (EMG) during wakefulness with slow respirations (left column), slow-wave sleep with slow respirations (middle column) and wakefulness with fast respirations (right column), all of which occurred in a single recording session. Upward deflections of the respiration trace indicate inhalation, while downward swings show exhalation. (B) Average of LFPs and respiration traces aligned in reference to exhalation onset (peak of the respiration trace) during wakefulness with slow respirations (left; average of 20s data during continuous slow respiration), slow-wave sleep with slow respirations (middle; average of 100s data during continuous slow respiration) and wakefulness with fast respirations (right; average of 4s data during continuous fast respiration) in a single recording session. Vertical broken line indicates the exhalation onset. (C) (left) Average power spectra of LFPs in the olfactory cortex during wakefulness with slow respirations (AON: green line, n = 9 sites from 4 rats; APC: magenta line, n = 11 sites from 5 rats; PPC: purple line, n = 10 sites from 3 rats) and that of respiration trace (blue bars, n = 30 sessions from 6 rats); (right) those during slow-wave sleep (AON: n = 4 sites from 3 rats; APC: n = 5 sites from 4 rats; PPC: 5 sites from 2 rats; respiration: 14 sessions from 4 rats). LFP powers are expressed as the arbitrary unit (a.u.) in this figure and other figures of this paper. Respiration powers are adjusted in each figure. Thin dotted lines, mean ± SEM. (D) Mean LFP powers of 0.5-3Hz band in each recording site during wakefulness with slow respirations (blue bars) and slow-wave sleep (gray bars). Welch’s t test; AON: Wake (n = 9) vs. Sleep (n = 4), T(8) = 0.98, p = 0.36; APC: Wake (n = 11) vs. Sleep (n = 5), T(4) = −2.34, p = 0.08; PPC: Wake (n = 10) vs. Sleep (n = 5), T(7) = −2.36, p = 0.05. Error bars, mean ± SEM. (E) Coherence between the LFPs and respiration during wakefulness with slow respirations (left) and slow-wave sleep (right). Thin dotted lines, mean ± SEM (F) Mean coherence between the LFPs and respiration trace in 0.5-3Hz band. Welch’s t test; AON: Wake (n = 9) vs. Sleep (n = 4), T(11) = 1.85, p = 0.09; APC: Wake (n = 11) vs. Sleep (n = 5), T(12) = 2.63, p = 0.02; PPC: Wake (n = 10) vs. Sleep (n = 5), T(13) = −3.57, p = 0.003). *p < 0.05. Error bars, mean ± SEM.

During awake state with slow respirations, LFPs in the deep layers of the AON (layer II), APC (layer III), and PPC (layer III) showed large-amplitude slow oscillatory potentials of 0.5-3Hz that were coherent with the slow respiration rhythm (Fig. 1 A-left and B-left in the APC; C-left and E-left in the AON, APC and PPC). These slow oscillatory potentials were similar to those reported in anesthetized rats (Fontanini et al., 2003: Fontanini and Bower, 2005). During awake state with fast respirations, the LFPs showed smaller-amplitude oscillations that were coherent with the fast respiration rhythms (3-7 Hz) (Fig. 1 A-right and B-right in the APC). Large-amplitude slow oscillatory LFPs were also observed in the olfactory cortical areas during slow-wave sleep state (Fig. 1 A-center and C-center).

The mean powers of the slow oscillatory potentials (0.5-3Hz) in the AON, APC and PPC during slow-wave sleep were not significantly different from those during awake state with slow respirations (Fig. 1D). However, the coherence between the slow oscillatory potentials and the respiration rhythm was substantially smaller during slow-wave sleep state than awake state with slow respirations (Fig. 1 B-center, E-right, and F). The diminished coherence between the slow oscillatory LFPs and the slow respiration rhythm during slow-wave sleep indicates that most of the slow oscillatory LFPs occur independently of respiration-phased olfactory sensory inputs during slow-wave sleep. In addition, this observation raised the possibility that the slow oscillatory LFPs during awake state with slow respirations might contain a component that occurs independently of respiration-phased olfactory sensory inputs.

### Respiration phases and layer-specific synaptic inputs in the olfactory cortex

To examine neural mechanisms underlying the generation of the slow oscillatory LFPs in the olfactory cortex, we measured the laminar profile of LFPs during awake state with slow respirations and compared the LFP traces with the time course of inhalation and exhalation. The translaminar LFPs in the AON (Fig. 2 C), APC (Fig. 2 D) and PPC (Fig. 3) showed slow oscillatory potentials that systematically changed in amplitude and polarity in relation to the layers of the olfactory cortex. The slow oscillatory potentials flipped over at the border between layers I and II in each area of the olfactory cortex, indicating that the slow oscillatory potentials were not due to volume conduction from elsewhere in the brain but were generated by neural activities within each area of the olfactory cortex.

**Figure 2.**
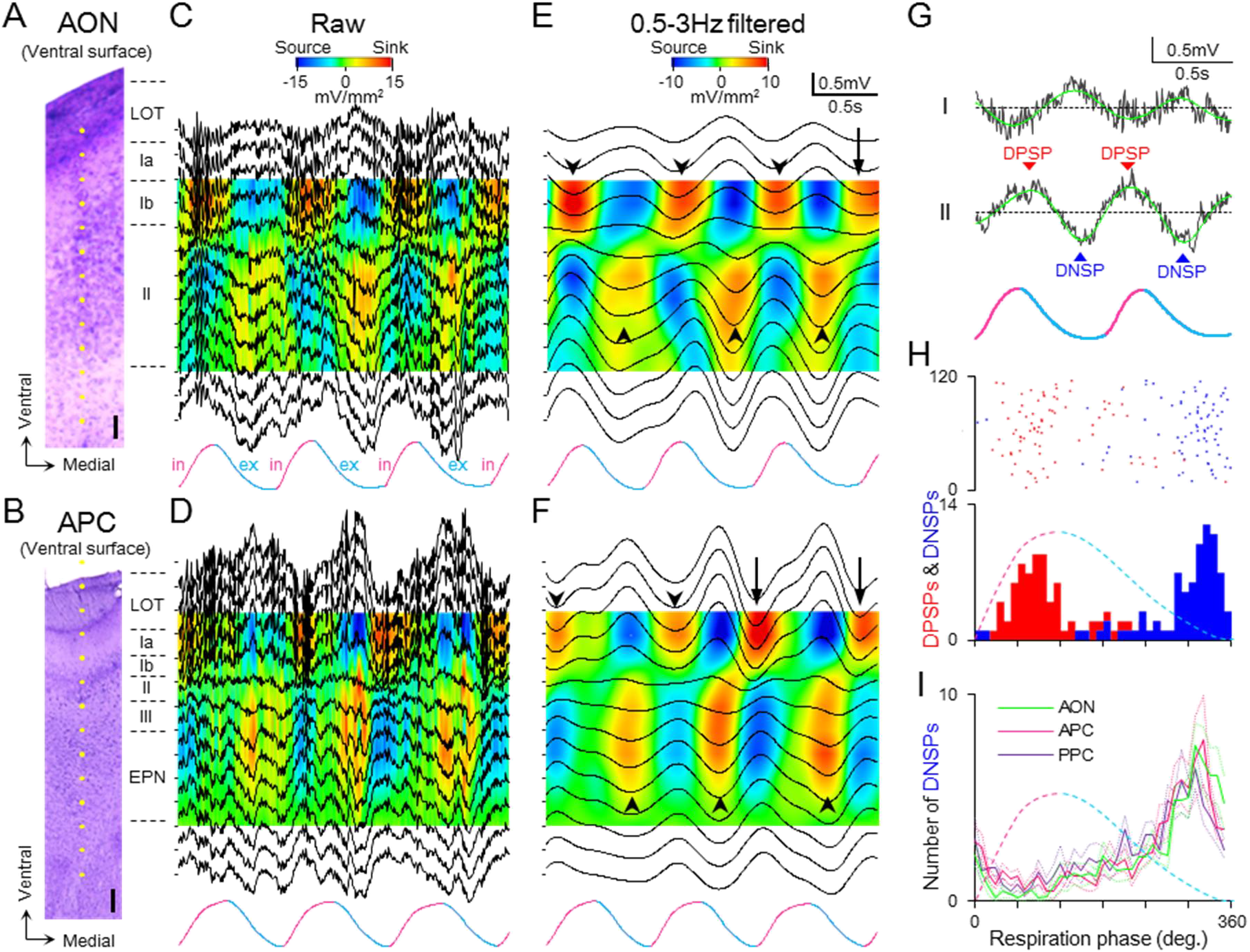
Superficial-layer current sinks during inhalation and deep-layer current sinks during exhalation. (A-F) Depths profiles and current source density (CSD) analysis of LFPs in the AON (A, C, E) and APC (B, D, F) with respiration trace (bottom). Inhalation phase (in) is indicated by magenta, while exhalation phase (ex) is indicated by blue on the respiration trace. In C and D, raw LFPs recorded at the estimated electrode positions (yellow dots in A and B, scale bar 100 μm) and the calculated CSD were aligned from the ventral surface (uppermost) to the depth. Warm colors in the CSD analysis indicate current sinks while cold colors show current sources. In E and F, the raw LFPs (C and D) were band-pass filtered (0.5-3 Hz) and then CSDs were calculated. Downward arrowheads indicate SL slow current sinks. Upward arrowheads indicate DL slow current sinks. Downward arrows indicate the SL slow current sinks whose onset preceded the inhalation onset. (G) An example of LFPs in the superficial layer (I) and deep layer (II) of the AON with respiration trace. Overlaid green trace is the 0.5-3Hz band-pass filtered LFP. Slow (0.5-3Hz) positive and negative potentials of deep layer LFP are defined here as depth-positive slow potential (DPSP, red arrowhead) and depth-negative slow potential (DNSP, blue arrowhead), respectively. (H) An example of the raster plots and the respiration phase histogram of the peak time of DPSP (red) and DNSP (blue) in the AON. Inhalation onset is at 0 degree and the end of exhalation at 360 degree. Light gray broken line is a putative respiration trace. (I) Average respiration phase histogram of the peak time of DNSP in the AON (green line, n = 9 sites from 4 rats), APC (magenta line, n = 12 sites from 5 rats), and PPC (purple line, n = 10 sites from 3 rats). DNSP in each area occurred predominantly at the latter part of the exhalation phase. Respiration phases of the peak times of AON DNSP, APC DNSP and PPC DNSP were 310, 320 and 310 degrees, respectively. All three phase histograms, p < 0.00001, Rayleigh test. Thin dotted lines, mean ± SEM EPN, endopiriform nucleus; LOT, lateral olfactory tract.

**Figure 3.**
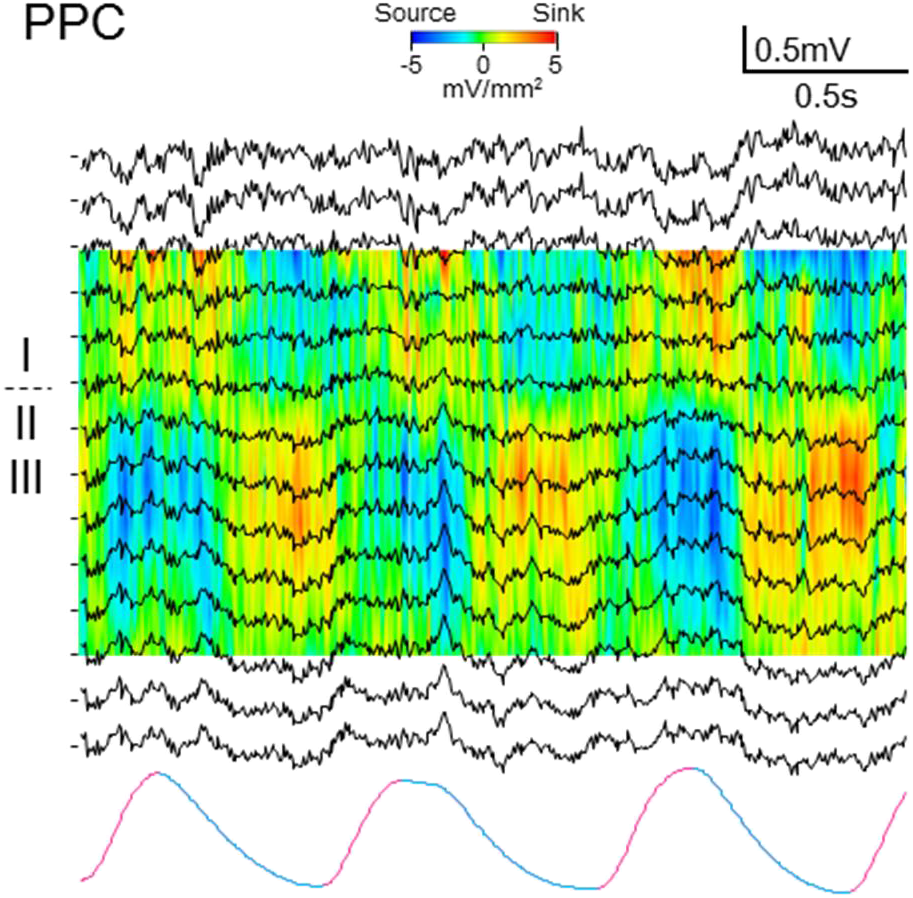
Depth profile of LFP and CSD in the PPC aligned with respiration trace. Inhalation phase is indicated by magenta colored respiration trace, while exhalation phase is indicated by blue (bottom trace). LFPs recorded at different depths and the calculated CSD were aligned from the ventral surface (uppermost) to the depth of PPC. Warm colors in the CSD indicate current sinks while cold colors show current sources.

CSD analysis of raw LFP traces revealed that a slow and long current sink in superficial layers (SL, layers Ia and Ib) (SL slow current sink) occurred in phase with inhalation while a slow and long current sink in deep layers (DL, layers II and III) (DL slow current sink) occurred in phase with exhalation (Fig. 2 C, AON; Fig. 2 D, APC; and Fig. 3, PPC). In phase with inhalation, the SL slow current sink was superimposed by a train of β- and γ-range (10-100Hz) fast oscillatory current sinks in the SL. In phase with exhalation, on the other hand, the DL slow current sink was overlapped with a train of β- and γ-range fast oscillatory current sinks in the DL (Fig. 2 C and D). Filtering out the β- and γ-range fast oscillatory current sinks (CSD analysis of 0.5-3Hz band-pass-filtered LFPs) clearly showed that the SL slow current sink (down-facing arrowheads and arrows in Fig. 2 E and F) occurred in synchrony with inhalation whereas the DL slow current sink (up-facing arrowheads in Fig. 2 E and F) occurred in synchrony with exhalation. In other words, the alternation between SL slow current sink and DL slow current sink occurred in synchrony with the alternation between inhalation and exhalation.

The SL-slow current sink typically started after inhalation onset and lasted for about 0.1-1.0 s (0.42 ± 0.14 s, mean ± SD, n = 331 SL-slow current sinks in the AON and APC) up to the early part of the subsequent exhalation phase. However, the onset of the SL slow current sink sometimes preceded the inhalation onset (arrows in Fig. 2 E and F), suggesting that the SL slow current sink can start spontaneously before the actual inhalation occurs. At the time window of the SL slow current sink, a slow current source in the deep layers (DL slow current source) was observed in each area. Thus, the AON, APC and PPC coordinately exhibited the SL slow current sink and DL slow current source during inhalation phase.

During the exhalation phase, on the other hand, the AON, APC and PPC showed a slow and long current sink in the deep-layers (DL slow current sink) (up-facing arrowheads in Fig. 2 E and F) and a slow and long current source in the superficial layers (SL slow current source). The exhalation-phased DL slow current sink lasted for about 0.1-1.0 s (0.41 ± 0.14 s, mean ± SD, n = 276 DL slow current sinks in the AON and APC). The end of exhalation-phased DL slow current sink typically coincided with the start of subsequent inhalation-phased SL slow current sink. Thus, all these olfactory cortical areas coordinately showed the SL slow current source and DL slow current sink during exhalation phase.

We further examined the temporal relationship between the slow oscillatory activity in the olfactory cortex and the inhalation-exhalation cycle by making respiration-phase histograms of the slow oscillatory LFPs recorded during awake state with slow respirations. Comparison of the CSD data with the depth profile of LFPs indicated that the SL slow current sink during inhalation corresponded with a slow negative potential in the SL and a slow positive potential in the DL (depth-positive slow potential; DPSP) (Fig. 2 E-G, Fig. 3, and Fig. 4). It also indicated that the DL slow current sink during the exhalation was associated with a slow negative potential in the DL (depth-negative slow potential; DNSP) and a slow positive potential in the SL (Figs 2 E-G, Fig. 3, and Fig. 4). We therefore used the DPSP as an indicator of SL slow current sink and the DNSP as an indicator of DL slow current sink.

**Figure 4.**
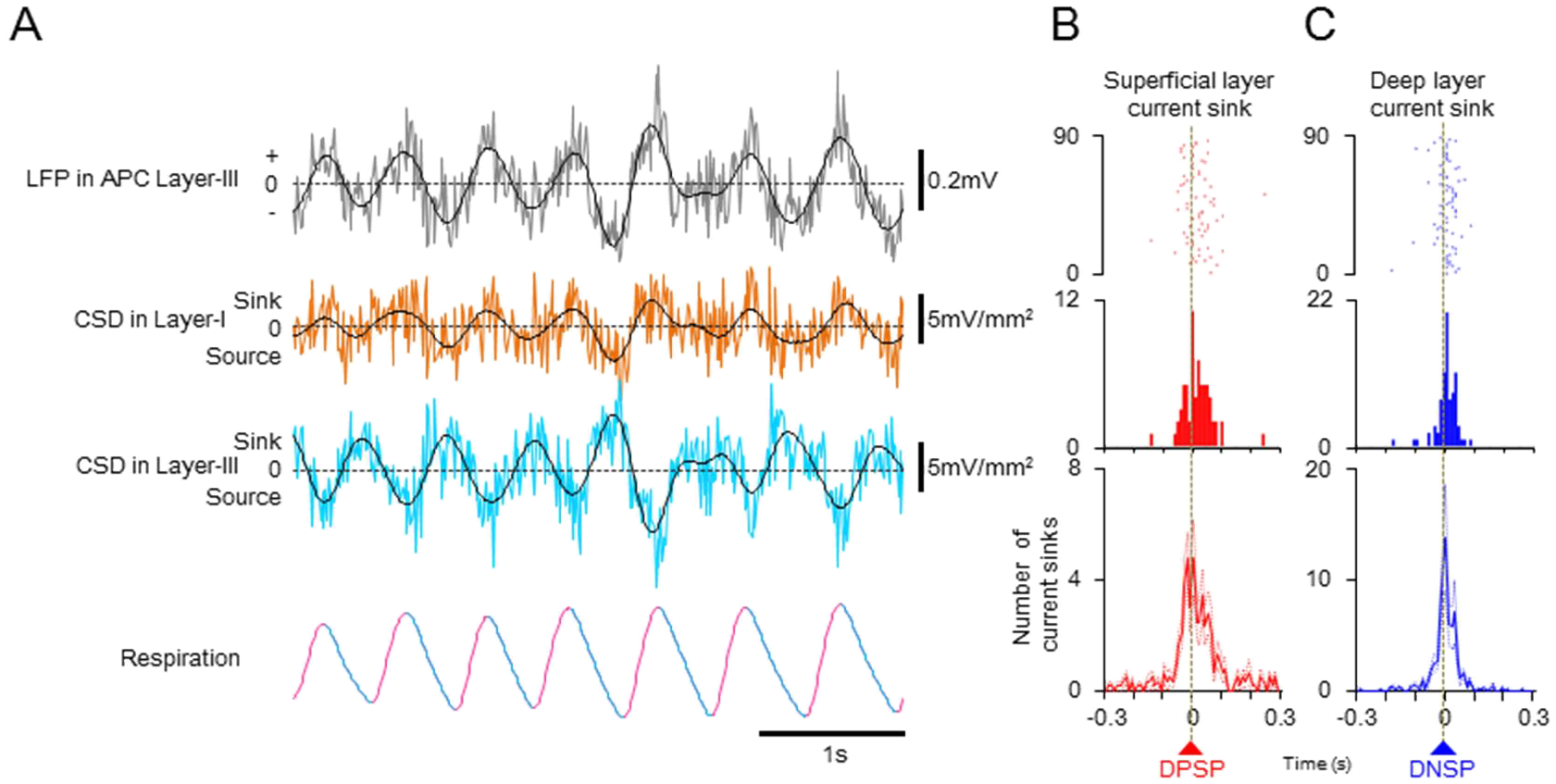
Depth-positive slow potential (DPSP) and depth-negative slow potential (DNSP) correlate tightly with superficial-layer slow current sink and deep-layer slow current sink, respectively. (A) An example of LFP and calculated CSDs in the APC. Top trace is the LFP recorded in the layer III of the APC. Second and third traces are CSD data in layer I and layer III respectively. Overlaid black line shows 0.5-3Hz band-pass-filtered LFP (top trace) or CSDs (second and third traces). Bottom trace represents respiration. (B, C) Perievent time histograms (PETH) of the peak of DPSP and the peak of slow current sink in the superficial layer (B), and PETH of the peak of DNSP and the peak of current sink in the deep layers (C). Peak time of each current sink in the superficial and deep layers was aligned to the peak time of DPSP and DNSP respectively. Top panels show examples of raster plot and PETH. Bottom panel showed average PETHs from 6 recordings in AON and APC. Thin dotted lines, mean ± SEM.

As exemplified in the respiration-phase histogram of DPSPs and DNSPs in the AON (Fig 2 H), the peak of DPSP (i.e., SL slow current sink) occurred mostly during inhalation phase, whereas the peak of DNSP (i.e., DL slow current sink) occurred predominantly during the latter half of exhalation phase. In all the areas (AON, APC and PPC), the peak of DPSP was phase-locked to the inhalation while the peak of DNSP was in phase with the latter half of exhalation, indicating that across the AON, APC and PPC, DPSPs occur in synchrony with inhalation while DNSPs occur in synchrony with the latter half of exhalation (Fig. 2 I, Fig. 4). These results indicate that the olfactory cortex areas coordinately show switching between SL slow current sink (DPSP) during inhalation and DL slow current sink (DNSP) during exhalation.

### Persistence of respiration-phased slow current sinks in ipsilateral naris-occluded rats

Are the inhalation-phased SL slow current sink and the exhalation-phased DL slow current sink in the olfactory cortex driven passively by inhalation-phased olfactory sensory inputs or generated actively and internally in the brain independent of inhalation-phased olfactory sensory inputs? To address this question, rats were subjected to ipsilateral naris occlusion with a nose plug. Because olfactory cortex receives olfactory sensory inputs mainly from ipsilateral olfactory epithelium via ipsilateral olfactory bulb, we expected that inhalation-phased olfactory sensory inputs would diminish under the condition of ipsilateral naris occlusion.

Inhalation of odor-containing air induces synchronized β- and γ-range fast oscillatory spike discharges of mitral and tufted cells in the olfactory bulb during the inhalation phase (Cury and Uchida, 2010; Manabe and Mori, 2013; Lepousez and Lledo, 2013). The sensory-driven fast oscillatory spike activities of olfactory bulb neurons are conveyed via their axons to the layer Ia of ipsilateral olfactory cortex (Fig. 8) (Neville and Haberly, 2004; Mori and Manabe, 2013). The sensory-driven fast oscillatory activities of bulbar neurons are also conveyed to layer Ib of the olfactory cortex via a relay in the AON and APC (Neville and Haberly, 2004).

In agreement with the olfactory sensory inputs via mitral and tufted cells, the ipsilateral naris occlusion diminished the inhalation-phased β- and γ-range (10-100Hz) fast oscillatory potentials in the AON and APC (Fig. 5 A and D). However, even under the condition of ipsilateral naris occlusion, we observed robust slow oscillatory potentials that were phase-locked to the respiration cycle (Fig. 5 A-right, E, F and G). The LFP power of 0.5-3Hz band did not differ significantly between the naris-open and ipsilateral naris-occluded conditions (Fig. 5 E) in contrast to that of β- and γ-range fast oscillatory potentials (Fig. 5 D), suggesting that the 0.5-3Hz slow oscillatory activity occurred independent of ipsilateral olfactory sensory inputs. Even under the condition of ipsilateral naris-occlusion, the coherence of the slow oscillations of LFPs and the respiration rhythm did not change significantly in all the three areas (Fig. 5 F), and peaks of DPSPs and DNSPs were still phase-locked to the inhalation and exhalation, respectively (Fig. 5 G). This observation suggests that the phase-locking between the slow oscillatory LFPs and slow respiration occurs in the absence of inhalation-phased ipsilateral olfactory sensory inputs.

**Figure 5.**
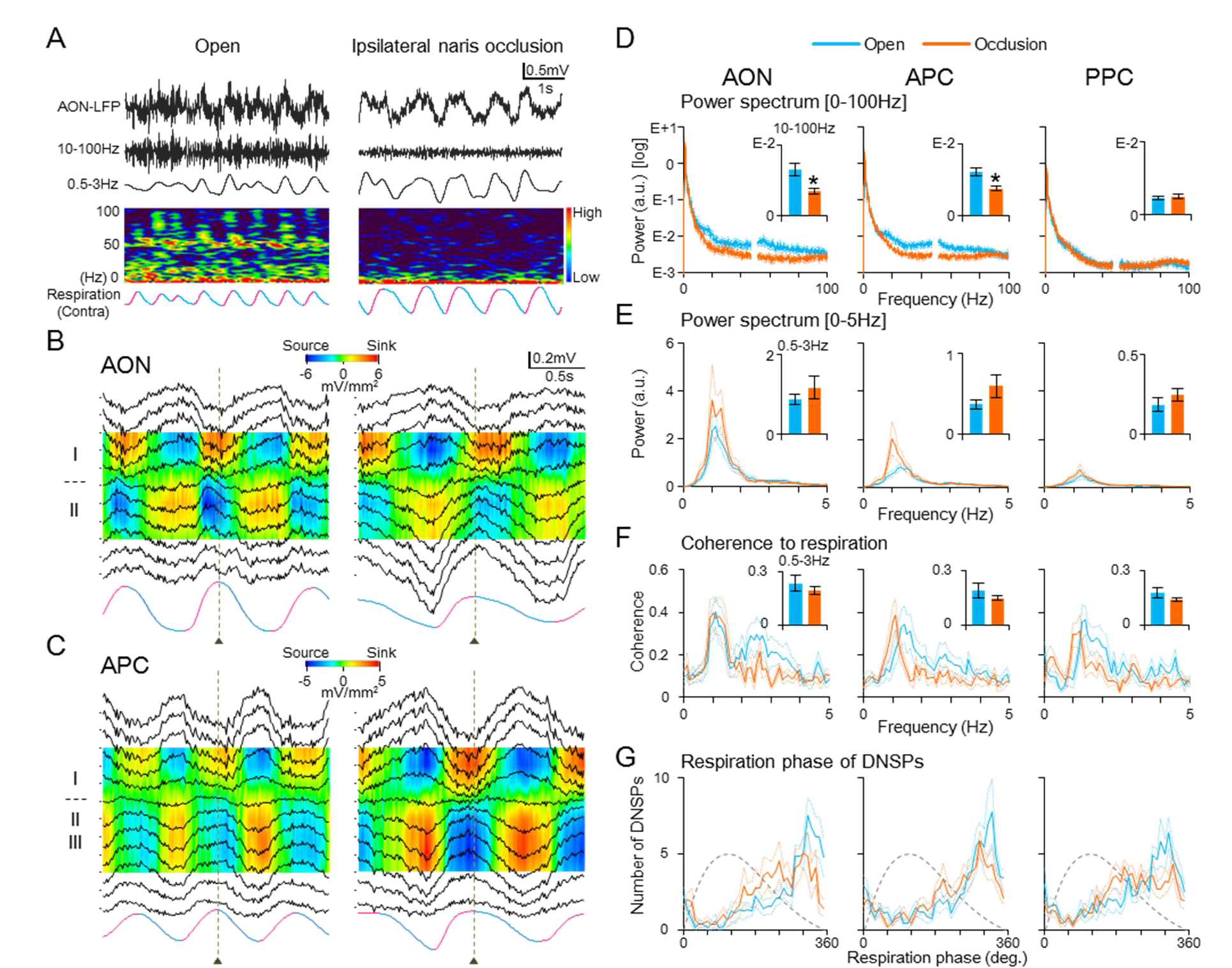
Effects of ipsilateral naris occlusion on respiration-phased slow oscillatory activity. (A) LFP in the deep layer of the AON under naris-open (left column) and ipsilateral naris-occluded (right column) conditions. The raw LFPs (uppermost trace) were band-pass filtered at 10-100Hz (2nd trace) or 0.5-3Hz (3rd trace). The colored maps show the time-frequency plots of the LFP power (0-100Hz). Respiration was monitored from the contralateral nostril (bottom trace). Note that high frequency signals (10-100Hz) are diminished by naris occlusion, whereas low frequency signals (0.5-3Hz) remained unchanged or was augmented. (B and C) Representative examples of depth profiles of LFP and their CSD in the AON (B) and APC (C) under the naris-open (left column) and naris-occluded (right column) conditions. The LFPs were averaged in reference to the onset of contralaterally-recorded exhalation (vertical broken line). Average respiration traces are shown at the bottom. (D-G) (left column) Data from AON, n = 9 sites from 4 rats; (middle column) data from APC, n = 11 sites from 5 rats; (right column) data from PPC, n = 10 sites from 3 rats. Blue lines and bars represent data in naris-open condition and orange lines and bars represent data in ipsilateral naris-occluded condition. Thin dotted lines and error bars, mean ± SEM (D) Average power spectra of the LFPs (0-100Hz range) in each area under the naris-open and naris-occluded conditions. Inserted bar graphs represent average LFP power of 10-100Hz band in each condition. Paired t test; AON: T(8) = 5.13, p = 0.0009; APC: T(10) = 7.39, p = 0.00002; PPC: T(9) = −1.103, p = 0.3. *p < 0.001. LFP power in high frequency range (10-100Hz) is significantly reduced in naris-occluded condition compared to naris-open condition in the AON and APC. (E) Magnification of low frequency range (0-5Hz) of the line graphs in D. Graphs of the naris-open condition are the same as those of wakefulness condition in Figure 1C and 1D. Inserted bar graphs represent average LFP power of 0.5-3Hz band in each condition. Paired t test; AON: T(8) = −1.4, p = 0.2; APC: T(10) = −1.78, p = 0.11; PPC: T(9) = −1.03, p = 0.33. Naris occlusion did not reduce the LFP powers of 0.5-3Hz band in all three areas. (F) Coherence between the LFPs and respiration trace under the naris-open and naris-occluded conditions. Graphs of the naris-open condition are the same as those of wakefulness condition in Figures 1E and 1F. Inserted bar graphs represent average coherence of 0.5-3Hz range in each condition. Paired t test; AON: T(8) = 1.31, p = 0.23; APC: T(10) = 1.28, p = 0.23; PPC: T(9) = −1.45, p = 0.18. Naris occlusion did not significantly reduce the coherence of 0.5-3Hz range in all three areas. (G) Average respiration phase histogram of DNSP peaks in each area under the naris-open and naris-occluded conditions. Line graphs of the naris-open condition are the same as those of Figure 2I. DNSP peaks in each area occurred at the latter part of exhalation phase even under ipsilateral naris-occluded condition. Respiration phases of the peak times of the AON DNSP, APC DNSP and PPC DNSP were 300, 290 and 320 degrees, respectively, in the naris-occluded condition. All three areas, p < 0.00001, Rayleigh test.

CSD analysis of LFPs averaged in reference to the exhalation onset revealed that even under the ipsilateral naris-occluded condition, SL slow current sink occurred during inhalation while DL slow current sink occurred during the exhalation in the AON (Fig. 5 B) and APC (Fig. 5 C), corroborating the idea that the SL slow current sink during inhalation and the DL slow current sink during exhalation occur in the olfactory cortex even without inhalation-phased ipsilateral olfactory sensory inputs.

### Exhalation-phased firing of olfactory cortex neurons in ipsilateral naris-occluded rats

The SL slow current sink together with DL slow current source during inhalation reflects larger slow depolarization of membrane potential in apical tuft dendrites (SL-dendrites) of pyramidal cells compared with that of proximal apical dendrites and basal dendrites (DL-dendrites). We asked the question whether the larger slow depolarization of SL-dendrites by itself can induce spike output of pyramidal cells or it has a modulatory role assisting the concomitant olfactory sensory inputs to the SL in generating spike outputs.

To address this question, we recorded extracellular spike activities of individual neurons in the olfactory cortex with or without ipsilateral naris occlusion. In the control naris-open condition, a subset of neurons in the olfactory cortex showed enhanced spike activity during inhalation (AON: 45% [40/88 cells], Figs. 6 A-left, B-left and C-upper left; APC: 34% [35/102 cells], D-left) and a different subset showed enhanced spike activity during exhalation (AON: 44% [39/88 cells], Figs. 6 B-left and C-lower left; APC: 40% [41/102 cells], D-left). The ipsilateral naris occlusion drastically decreased the inhalation-phased spike activities both in the AON (to 13% [11/88 cells]) and APC (to 13% [13/102 cells]) (Figs. 6 B-D), suggesting that the enhanced spike activities during inhalation under the naris-open condition are driven mainly by inhalation-phased olfactory sensory inputs. These results suggest that the slow and long depolarization of apical tuft dendrites in SL during inhalation does not induce spike outputs by itself but has a modulatory role assisting the concomitant olfactory sensory inputs to induce spike outputs.

**Figure 6.**
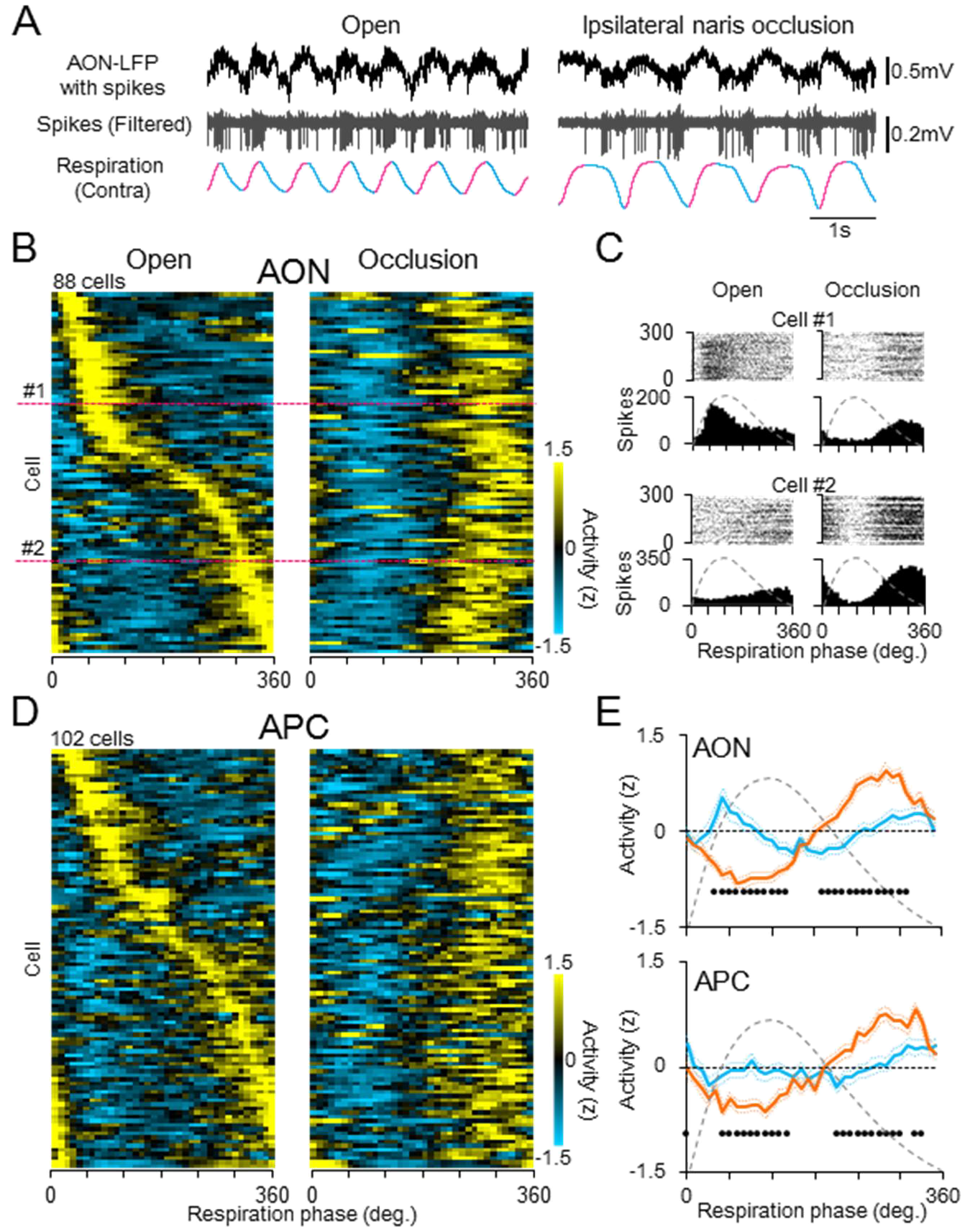
Effects of ipsilateral naris occlusion on spike timing of the olfactory cortical neurons. (A) LFP and single-cell spike activities recorded in the AON under naris-open (left) and ipsilateral naris-occluded (right) conditions in a single recording session. Spike activity of this neuron during inhalation was diminished by naris occlusion, whereas that during exhalation remained unchanged or was augmented. (B) Population data of the respiration phase histograms of spike activities of individual neurons in the AON (88 cells from 3 rats) under naris-open (left) and ipsilateral naris-occluded (right) conditions. Spike activities are expressed as z-score of each neuron and coded in colors (high: yellow; 0: black; low: blue). Cells were sorted by the respiration phase of the peak spike activity in the naris-open condition. Cells #1 and #2 (magenta broken lines) in population data correspond to those in C. (C) Raster plots and a respiration phase time histogram of spike activity of two representative neurons (Cells #1 and #2) in the AON under naris-open (left column) and ipsilateral naris-occluded (right column) conditions. (D) Population data of the respiration phase histograms of spike activities of individual neurons in the APC (102 cells from 4 rats) under naris-open (left) and ipsilateral naris-occluded (right) conditions. The same format as B. (E) Average data of the respiration phase time histogram of spike activity under naris-open (blue trace) and ipsilateral naris-occluded (orange trace) conditions in the AON (top) and APC (bottom). Significant differences between groups are represented as black dots in each degree (p < 0.01, paired t-test). Thin dotted lines, mean ± SEM

The DL slow current sink together with SL slow current source during exhalation reflects larger slow depolarization of membrane potential in proximal apical dendrites and basal dendrites in DL compared with that in apical tuft dendrites in the SL. We found that the ipsilateral naris-occlusion markedly increased the number of AON neurons (80% [70/88 cells], Fig. 6 B-right) and APC neurons (89% [78/88 cells], Fig. 6 D-right) that showed enhanced spike activities during exhalation. The results indicate that even in the absence of inhalation-phased ipsilateral olfactory sensory inputs, the brain generates exhalation-phased slow depolarization in DL dendrites of olfactory cortex pyramidal cells so that a majority of olfactory cortex neurons show enhanced spike activities during exhalation.

### Persistence of exhalation-phased slow current sink with spike activities after bilateral olfactory bulb removal

Because the olfactory cortex also receives contralateral olfactory sensory inputs through association fibers crossing the anterior commissure (Haberly and Price, 1978; Luskin and Price, 1983), the ipsilateral naris occlusion cannot eliminate the effect of inhalation-phased sensory inputs from the contralateral olfactory epithelium. To address this possibility, we surgically ablated both the right and left olfactory bulbs (bilateral olfactory bulbectomy: OBX) so that the olfactory sensory inputs are completely eliminated. Respiration-phased neural activities were examined in the olfactory cortex of these bilaterally bulbectomized rats.

In all the three areas of OBX rats, inhalation-phased β- and γ-range fast oscillations of LFPs were greatly reduced compared to naris-open intact group or even compared to ipsilateral naris-occluded group (Fig. 7 A), indicating that inhalation-phased olfactory sensory inputs are successfully blocked by OBX. On the other hand, the power of LFP slow oscillations (0.5-3Hz) did not change in the APC and PPC (Fig. 7 B), confirming that the slow oscillations in the APC and PPC occur independent of respiration-phased olfactory sensory inputs. In the AON of OBX rats, however, we observed a significant decrease of slow oscillatory potentials (0.5-3Hz) (Fig. 7 B), implicating that that the slow oscillation in the AON consists of two components; one component occurs independently of olfactory sensory inputs and the other is generated by the summation of fast synaptic potentials of inhalation-phased olfactory sensory inputs.

**Figure 7.**
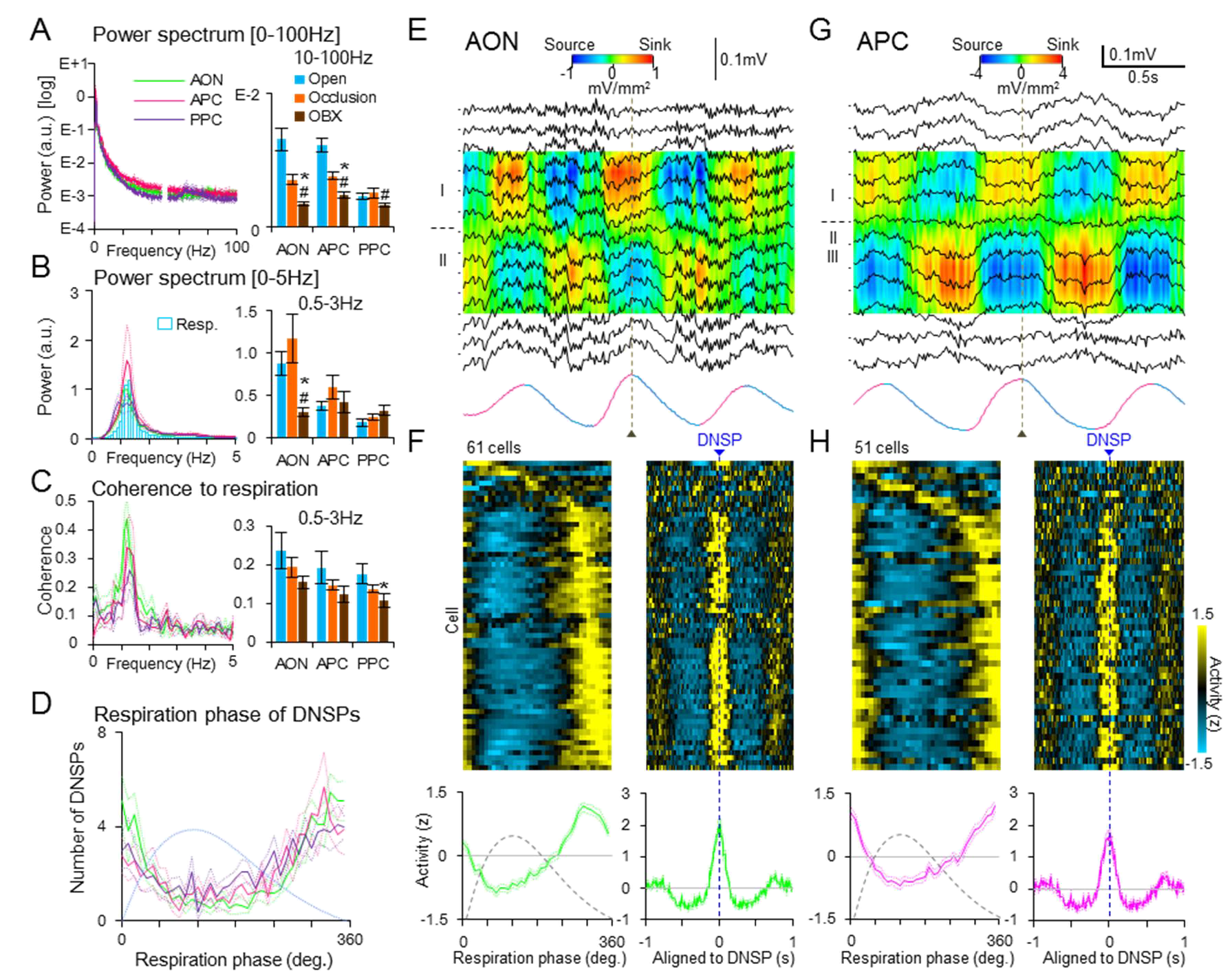
Local field potential and spike activities of olfactory cortex in bilaterally bulbectomized rats. (A-D) Data from 4 bilaterally bulbectomized (OBX) rats: AON, n = 11 sites; APC, n = 9 sites; PPC, n =10 sites. Green, magenta and purple lines represent average data of AON, APC and PPC, respectively. Thin dotted lines, mean ± SEM. In A-C, bar graphs represent average data of certain frequency ranges in OBX (brown bars), naris-open (Open, blue bars) and ipsilateral naris-occluded (Occlusion, orange bars) rats. Bar graphs of Open and Occlusion group are redrawing of those in Fig. 5 D-F for comparisons. Statistical test was conducted within each area using Welch’s t test against OBX. *p < 0.05, OBX vs. Open; #p < 0.05, OBX vs. Occlusion. Error bars, mean ± SEM (A) (left) Average power spectra of the LFPs (0-100Hz range) in the AON, APC and PPC of OBX rats. (right) Bar graphs represent average LFP power of high frequency (10-100Hz) range in each area. AON: OBX vs. Open, T(9) = 5.66, p = 0.0003; OBX vs. Occlusion, T(10) = 4.18, p = 0.002; APC: OBX vs. Open, T(14) = 6.34, p = 0.00002; OBX vs. Occlusion, T(18) = 3.88, p = 0.001; PPC: OBX vs. Open, T(14) = 2.5, p = 0.03; OBX vs. Occlusion, T(11) = 2.42, p = 0.03. The LFP power of the high frequency range is reduced in all three areas of OBX group compared to Open and Occlusion groups. (B) (left) Magnification of low frequency range (0-5Hz) of the line graphs in A. Opened bar graph represent mean power spectrum of respiration trace in OBX rats. (right) Bar graphs represent average LFP power of 0.5-3Hz band in each area. AON: OBX vs. Open, T(10) = 3.85, p = 0.003; OBX vs. Occlusion, T(9) = 2.99, p = 0.02; APC: OBX vs. Open, T(11) = −0.35, p = 0.74; OBX vs. Occlusion, T(18) = 0.91, p = 0.37; PPC: OBX vs. Open, T(17) = −1.96, p = 0.07; OBX vs. Occlusion, T(16) = −1.11, p = 0.28. The LFP power of this range was significantly reduced in the AON compared to Open and Occlusion groups, but not in the APC and PPC. (C) (left) Coherence between the LFPs and respiration trace of OBX group. (right) Bar graphs represent average coherence of 0.5-3Hz range in each area. AON: OBX vs. Open, T(10) = 1.64, p = 0.13; OBX vs. Occlusion, T(14) = 1.31, p = 0.21; APC: OBX vs. Open, T(14) = 1.46, p = 0.17; OBX vs. Occlusion, T(14) = 0.98, p = 0.34; PPC: OBX vs. Open, T(15) = 2.21, p = 0.04; OBX vs. Occlusion, T(15) = 1.61, p = 0.13. The coherence of this range was slightly reduced in the PPC, but no significant reduction was observed in the AON and APC. (D) Average respiration phase histogram of DNSP peaks in each area of OBX group. DNSP peaks in each area occurred at the latter part of exhalation phase even under OBX condition. Respiration phase of the peak times of AON DNSP, APC DNSP and PPC DNSP were 330, 320 and 340 degrees, respectively. All phase histograms, p < 0.00001, Rayleigh test. (E, G) Depth profiles of LFPs and their CSD analysis in the AON (E) and APC (G) under OBX condition. The LFPs and respirations were averaged in reference to exhalation onset (vertical broken line). (F, H) (Top panels) Population data of spike activities of individual neurons in the AON (F, 61 cells from 3 OBX rats) and APC (H, 51 cells from 3 OBX rats) in reference to respiration cycle (respiration phase histogram, left) or to the peak of DNSP (perievent time histogram, right) of the bulbectomized rats. Cells were sorted by the respiration phase of the peak spike activity in the respiration phase histogram. (Bottom panels) Average data of the respiration phase time histogram and perievent time histogram of spike activities.

The slow oscillatory LFPs in the AON, APC and PPC showed clear coherence with respiration rhythm in OBX rats (Fig. 7 C). In addition, DNSPs of all the three areas were phase-locked to the exhalation to the similar extent as in control rats (Fig. 7D). These results are consistent with those in ipsilateral naris-occluded rats (Figs. 5 E-G) and again support the idea that the slow oscillatory activities in the olfactory cortex areas occur in synchrony with the respiration rhythm even under the condition of total absence of inhalation-phased olfactory sensory inputs.

CSD analysis of the LFPs in the AON and APC of OBX rats showed the alternation between the inhalation-phased SL slow current sink (DL slow current source) and the exhalation-phased DL slow current sink (SL slow current source) (Figs. 7 E and G) as is the case in control open naris rats. Unit recordings revealed that spike activities of a majority of neurons in both the AON (Fig. 7 F-left) and APC (Fig. 7 H-left) were greatly reduced during inhalation, suggesting that, without olfactory sensory input, most of neurons in the AON and APC are nearly silent during inhalation.

In a striking contrast, a majority of neurons in the AON (89% [54/61 cells], Fig. 7 F-left) and APC (61% [31/51 cells], Fig. 7 H-left) of OBX rats showed enhanced spike activity during exhalation. Exhalation-phased DL slow current sink (SL slow current source) accompanying with a large number of exhalation-phased spike activities in OBX rats suggests that the DL slow current sink is generated by slow depolarization of basal dendrites of pyramidal cells and that the slow depolarization facilitates top-down inputs to generate spike outputs. Consistent with this notion, spike activities of the AON neurons and APC neurons were in synchrony with the peak of DNSP that was associated with the exhalation-phased DL slow current sink (Fig. 7 F-right and H-right). These results from OBX rats support the idea that exhalation-phased slow depolarization of DL dendrites and exhalation-phased spike activities of olfactory cortex neurons occur even in total absence of inhalation-phased olfactory sensory inputs.

## Discussion

### (1) Inhalation-phased SL slow current sink and exhalation-phased DL slow current sink in the olfactory cortex

In the present study, the CSD analysis of LFPs revealed that when rats were in awake state with slow respirations, the AON, APC and PPC showed two distinct types of current sinks in the SL that occurred in phase with inhalation: β- and γ-range fast oscillatory current sinks and a slow current sink (SL slow current sink) that typically lasted for about 300-550 msec.

Because odor inhalation causes β- and γ-range fast oscillatory spike activities of mitral and tufted cells in the olfactory bulb that send axons and terminate on apical tuft dendrites of pyramidal cells in layer Ia of the olfactory cortex (Neville and Haberly, 2004), mitral and tufted cell axons may convey the inhalation-phased olfactory sensory signals to the olfactory cortex and generate the β- and γ-range fast oscillatory current sinks in the SL during inhalation (Fig. 8). Since pyramidal cells in the AON give rise to Ib association fibers to pyramidal cells of AON, APC and PPC (Neville and Haberly, 2004), synaptic inputs from the Ib association fibers might also participate in the generation of the β- and γ-range fast oscillatory current sinks in the SL during inhalation.

**Figure 8.**
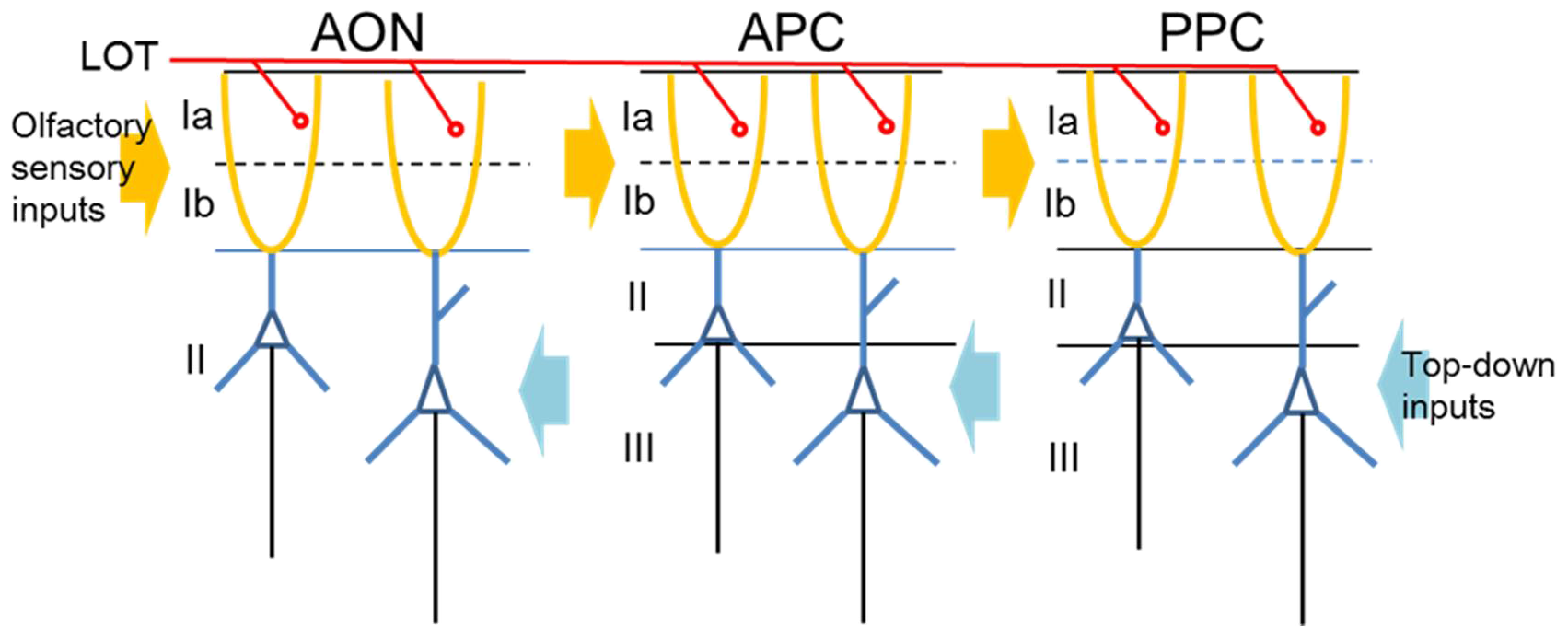
A schematic diagram illustrating pyramidal cells with apical tuft dendrites (orange) in the superficial layers (SL, layers Ia and Ib) and proximal apical dendrites and basal dendrites (blue) in the deep layers (DL, layers II and III) in the anterior olfactory nucleus (AON), anterior piriform cortex (APC), and posterior piriform cortex (PPC). A slow depolarization occurs in apical tuft dendrites in the SL during inhalation whereas a slow depolarization occurs in proximal apical dendrites and deep dendrites in the DL during exhalation. Olfactory sensory inputs to the SL are shown by orange arrows, while top-down inputs to the DL by blue arrows. Red lines indicate axons of mitral and tufted cells of the olfactory bulb running through the lateral olfactory tract (LOT) and terminating in layer Ia.

In agreement with the idea that bottom-up olfactory sensory inputs cause the fast oscillatory current sinks in SL of the olfactory cortex areas, elimination of inhalation-phased olfactory sensory inputs either by ipsilateral naris occlusion (Fig. 5) or by bilateral bulbectomy (Fig. 7) diminished inhalation-phased β- and γ-range fast oscillatory current sinks in the SL.

Supprisingly, neither ipsilateral naris occlusion nor bilateral bulbectomy reduced the inhalation-phased SL slow current sink (Fig. 5), indicating that the inhalation-phased SL slow current sink was not driven by inhalation-phased bottom-up olfactory sensory inputs. In other words, the brain spontaneously generates the inhalation-phased SL slow current sink in the olfactory cortex even in the absence of inhalation timing information from the inhalation-phased olfactory sensory inputs. This idea is consistent with our observation that the SL slow current sink sometimes preceded the onset of inhalation (Fig. 2 C-F, arrows). These results indicate that neural mechanism for generating the inhalation-phased SL slow current sink in the olfactory cortex differs completely from that for generating inhalation-phased SL fast oscillatory current sinks (the bottom-up olfactory sensory inputs).

CSD analysis also revealed two types of current sinks in the DL that occurred in phase with exhalation: β- and γ-range fast oscillatory current sinks and a slow current sink in the DL (DL slow current sink) that lasted for about 300-550 msec. Because synaptic inputs to the DL of olfactory cortex originate mainly from orbitofrontal cortex, ventral agranular insular cortex, medial prefrontal cortex, amygdala and higher areas of the olfactory cortex (Datiche and Cattarelli, 1996; Illig, 2005; Neville and Haberly, 2004), the exhalation-phased β- and γ-range fast oscillatory current sinks in the DL presumably reflect top-down synaptic inputs from these higher order areas. Both ipsilateral naris occlusion and bilateral olfactory bulbectomy reduced the exhalation-phased β- and γ-range fast oscillatory potentials in the DL, suggesting that inhalation-phased olfactory sensory inputs affect the generation of top-down signals in these higher order areas.

In a striking contrast, neither ipsilateral naris occlusion nor bilateral bulbectomy reduced the exhalation-phased DL slow current sink, indicating that DL slow current sink occurred spontaneously, independent of inhalation-phased bottom-up olfactory sensory inputs. These results suggest that neural mechanism for generating the exhalation-phased DL slow current sink in the olfactory cortex differs from that for generating exhalation-phased DL fast oscillatory current sinks.

### (2) Possible functions and neural mechanisms of inhalation-phased SL slow current sink and exhalation-phased DL slow current sink

The inhalation-phased SL slow current sink always accompanied DL slow current source. Current flow in pyramidal cells during the inhalation-phased SL slow current sink and DL slow current source indicates that a larger depolarized membrane potential occurs in apical tuft dendrites (SL dendrites) of pyramidal cells compared with proximal apical dendrites and basal dendrites (DL dendrites). The more depolarization in the SL dendrites during inhalation may greatly enhance the spike response of pyramidal cell to inhalation-phased olfactory sensory inputs to the SL dendrites, while the less depolarization in the DL dendrites during inhalation may be less effective in enhancing spike response to top-down inputs to the DL dendrites.

Ipsilateral naris occlusion and OBX experiments showed that in the absence of olfactory sensory inputs, inhalation-phased spike activities of olfactory cortex neurons diminished, suggesting that the slow and long depolarization itself may rarely induce spike outputs of pyramidal cells, but the combination of the slow depolarization with olfactory sensory inputs may effectively induce spike outputs. Thus we hypothesize that the inhalation-phased slow-current sink plays a modulatory role biasing for olfactory sensory inputs to the SL dendrites so that they become more effective in inducing spike outputs.

The present study does not address the question of what neural mechanism generates the inhalation-phased SL slow current sink (slow depolarization of apical tuft dendrites) in the AON, APC and PPC. It has been reported in the mouse auditory cortex that foot-shock-induced activation of cholinergic neurons causes long lasting disinhibition (depolarization) of pyramidal cells, and that convergence of the long lasting disinhibition and auditory sensory input is necessary for associative fear learning (Letzkus et al., 2011). In the mouse primary visual cortex, locomotion causes neuromodulation-mediated disinhibition (depolarization) of pyramidal cells independent of visual stimulation and this modulatory disinhibition enhances the pyramidal cells’ response to visual sensory input (Polack et al., 2013; Fu et al., 2014). We thus speculate that similar disinhibitory mechanism might be a candidate for causing the inhalation-phased slow depolarization (SL slow current sink) in apical tuft dendrites of olfactory cortex pyramidal cells. In support of this idea, apical tuft dendrites of pyramidal cells of the olfactory cortex receive not only excitatory synaptic inputs conveying olfactory sensory signals but also inhibitory inputs from GABAergic interneurons in the SL (Suzuki and Bekkers, 2007, 2012; Kay and Brunjes 2014; Neville and Haberly 2004).

It has been reported that pupil fluctuation occurs in association with variation of cortical states during wakefulness, rapid dilations of the pupil reflect changes in adrenergic and cholinergic activities (Reimer et al., 2014, 2016), and pupil diameter fluctuates in association with respiratory cycle during awake resting state (Nakamura et al., 2018). We thus speculate that fluctuation of noradrenergic or cholinergic input to the olfactory cortex might occur in association with inhalation and exhalation. In fact, we found that a subset of neurons in the basal forebrain cholinergic nuclei showed increased firing during inhalation and decreased firing during exhalation (Narikiyo et al., unpublished observation). Therefore, it is likely that increased activity of cholinergic or noradrenergic inputs during inhalation might activate layer I inhibitory interneurons in the olfactory cortex and di-synaptically inhibit other types of inhibitory interneurons, resulting in the slow disinhibition (depolarization) of apical tuft dendrites (SL dendrites) of olfactory cortex pyramidal cells during inhalation. Further studies are necessary to examine this possibility and to understand neural mechanisms responsible for the active generation of inhalation-phased slow depolarization of SL dendrites and exhalation-phased slow depolarization of DL dendrites.

How do olfactory cortex pyramidal cells generate the slow depolarization of SL dendrites at the timing of inhalation and the slow depolarization of DL dendrites at the timing of exhalation even in the absence of the inhalation/exhalation timing information from inhalation-phased olfactory sensory inputs? One possibility is that respiration-phased activity of trigeminal somatosensory afferents from the airway that remains after the bilateral OBX or the efference copy from the respiratory motor centers may be responsible for the generation of respiration-phased slow depolarizations. However this idea is inconsistent with the observation that the SL slow current sink sometimes occurred prior to the start of inhalation.

Another intriguing possibility is that when necessary the slow rhythm of respiration is generated intrinsically in the prefrontal cortex. Based on the previous reports describing that the insular cortex and the medial prefrontal cortex can modulate the respiration pattern (Dutschmann and Dick, 2012; Hassan et al, 2013), it is possible that the prefrontal cortices might generate the slow rhythm and send the signals both to the respiratory central pattern generator in the brainstem and to neural substrates that cause inhalation-phased slow depolarization of SL dendrites and exhalation-phased slow depolarization of DL dendrites of olfactory cortex pyramidal cells. In support of this hypothesis, it has been reported that the medial prefrontal cortex shows LFPs and spike activities that occur synchronized with nasal respiration rhythm during awake immobility (Biskamp et al., 2017). Further studies are necessary to examine these possibilities.

### (3) Respiration-phased switching between sensory inputs and top-down inputs

Individual pyramidal cells of the olfactory cortex are thought to function as coincidence detectors integrating synaptic inputs conveying olfactory sensory signals impinging on apical tuft dendrites and top-down synaptic inputs of contextual signals on proximal apical dendrites and basal dendrites (Shiotani et al., 2018, Sakmann, 2017; Hill et al., 2013). The present study indicates that the integration occurs in different manners during inhalation and exhalation: during inhalation, SL dendrites receive slow modulatory depolarizing inputs presumably to boost olfactory sensory inputs to be more effective in inducing spike outputs compared with top-down inputs, whereas during exhalation, slow modulatory depolarization occurs in DL dendrites biasing for top-down contextual signals (Fig. 8).

Based on these findings, we propose the following hypothesis of cortical mechanism to alternate attention either to external odor world or to internal contextual signals (Chun et al., 2011) in synchrony with the change in the direction of respiration, i.e., inhalation and exhalation. When prefrontal cognitive processes try to attend to external odor world, they presumably send top-down command signals to respiratory central pattern generator in the brainstem (Dutschmann and Dick, 2012) to cause active inhalation of odors. At the same time, they may activate neural systems that induce slow depolarization in apical tuft dendrites (SL dendrites) of olfactory cortex pyramidal cells, which enhances the pyramidal cells’ response selectively to inhalation-phased olfactory sensory inputs arriving in the SL (Letzkus et al., 2013; Polak et al., 2013). In this way, attention to external odor world may be coupled with active inhalation.

On the contrary, when prefrontal cognitive processes try to attend to internal contextual information, they presumably send top-down command signals to the brainstem respiratory central pattern generator to start active exhalation. At the same time, they may send signals to neural systems that induce slow depolarization in proximal apical dendrites and basal dendrites (DL dendrites) of olfactory cortex pyramidal cells, which facilitate the pyramidal cells’ response selectively to top-down contextual signals such as top-down scene signals (Letzkus et al., 2011; Polack et al., 2013; Shiotani et al., 2018) coming to the DL. Thus, attention to the internal contextual signals may be coupled with exhalation.

Further studies are needed to test the hypothesis of respiration-phased alternation of external attention and internal attention, and to understand the functional roles of the inhalation-phased slow depolarization of SL dendrites and exhalation-phased slow depolarization of DL dendrites in the olfactory cortex.

## Methods

### Animals

Male adult Long Evans rats (260-370 g) were obtained from Japan SLC. Animals were maintained with ad libitum food and water in a 12:12 hour light/dark cycle (Light phase: 5:00-17:00). All experiments were performed in accordance with the guidelines of the Physiological Society of Japan and were approved by the Experimental Animal Research Committee of the University of Tokyo.

### Surgery

Rats were implanted with recording electrodes under anesthesia with ketamine (60 mg/kg i.p.) and medetomidine (0.5 g/kg i.p.). During the surgery, body temperature was maintained at 37°C using a heat pad. An incision was made on the scalp with a subcutaneous injection of 0.1% lidocaine and skull surface was exposed. For electrical stimulation and LFP recording, three Teflon-insulated wires (75 μm diameter, A-M systems) were twisted, in which two wires were used as a bipolar stimulation electrode and the remaining wire as a recording electrode. The twisted electrodes were stereotaxically implanted into the olfactory bulb (OB; AP: 7.5 mm, ML: 1.2 mm, DV: 2.5 mm), anterior olfactory nucleus (AON; AP: 5.0 mm, ML: 2.0 mm, DV: 4.0 mm) or anterior piriform cortex (APC; AP: 2.0 mm, ML: 3.5 mm, DV: 6.5 mm) and fixed with dental acrylic.

For electrical ground and reference electrode, two stainless-steel screws were threaded into the bone above the cerebellum. For electrocorticogram (ECoG), another screw was threaded into the bone above the occipital cortex (AP: −6.3 mm, ML: 3.0 mm). Electromyogram (EMG) was recorded from neck muscle by a wire electrode. All electrodes were connected with electrical interface board (Neuralynx).

For later recording with silicon probes, cranial windows were made on the skull. The bone above the target sites of AON (AP: 2.0-5.5 mm, ML: 0.0-3.5 mm), APC (AP: 0.0-3.5 mm, ML: 2.0-5.5 mm) or posterior piriform cortex (PPC; AP: −1.0–4.5 mm, ML: 2.0-6.0 mm) was removed to make a small hole (typically 3-4 mm square area) by dental drill, leaving the dura matter intact. The small holes were covered with silicone impression material (Shofu) until the recording day. Finally, a head holder was fixed to the bone with anchor screws and adhesives.

For olfactory sensory deprivation experiments, four rats received bilateral olfactory bulbectomy. Under the anesthesia with ketamine (60 mg/kg i.p.) and medetomidine (0.5 g/kg i.p.), the bone above the olfactory bulb was removed by a dental drill and bilateral olfactory bulbs were surgically removed. Remaining olfactory nerves on the cribriform plate were carefully removed using fine forceps. The empty space was filled with dental silicone to prevent reinnervation of olfactory sensory neurons.

### Recordings

Neural activities were recorded using the chronically implanted electrodes and the acutely inserted 32ch silicon probe with 50 μm spacing linear contact sites (Impedance: 0.5-3 MΩ at 1 kHz) (Neuronexus). Electrical signals were filtered and sampled (for LFP: 0.1-6000 Hz filtering and 2 k or 15 kHz sampling; for spike activity: 600-6000 Hz filtering and 30 kHz sampling) using 48ch recording system (Neuralynx). To monitor respiration pattern, a thin wire thermocouple (0.2 mm diameter, World Precision Instruments) was inserted into the nostril (contralateral to the recording olfactory cortex), and the respiration-coupled temperature change in the nasal cavity was continuously measured by electronic digital thermometers (Physitemp).

After 1 week recovery period, animals had acclimatization to head-fixing experimental procedures. The head of animal was fixed to the stereotaxis apparatus (Narishige) using the previously implanted head holder. Trunk of animal was mildly restrained by covering with cylinder. To facilitate the habituation, 0.1% saccharin solution was supplied by a tube to the mouth of the animal in the head-fixed position. Animals also had acclimatized for inserting thermocouple and nasal plug into the nostrils. Animals adapted these experimental conditions within several days and showed quiet wakefulness state or spontaneous sleep for the most of the time during head-fixation.

On the recording day, silicone material on the cranial hole was gently removed, and dura was exposed and locally anesthetized with 1% lidocaine solution. DiI solution (Life technologies) was applied backside of the silicon probe for marking and identifying recording tracks. The silicone probe was inserted through the dura into the target area of the olfactory cortex; AON (AP: 4.0-5.2 mm, ML: 1.0-3.5 mm, DV: 5.0-7.5 mm), APC (AP: 0.0-2.5 mm, ML: 2.5-4.5 mm, DV: 6.0-9.5 mm) or PPC (AP: −2.0–3.5 mm, ML: 5.0-6.5 mm, DV: 9.0-11.0 mm) according to the rat brain atlas (Paxinos and Watson, 1998). The silicone probe was tilted up to 30° on coronal plane to insert vertically to layers of each olfactory cortex area and lowered slowly into the target depth using a manipulator (Narishige). The depth of silicon probe in the olfactory cortex was determined by monitoring LFPs evoked by electrical stimulation of the olfactory bulb, AON or APC. After inserting the silicon probe, the cranial window was filled with saline to avoid drying of the surface.

For investigation of spontaneous activity patterns of the olfactory cortical areas, animals breathed freely the air of the experimental room and we did not apply any specific odors during experiments. To examine the effect of olfactory sensory input on the olfactory cortex activity, unilateral nostril (ipsilateral to recording hemisphere) was occluded temporarily by the nasal plug (rolled double-sided tissue tape). The nasal plug successfully diminished the respiration-coupled fast oscillations in the olfactory cortex, an indicator of olfactory sensory input (Fig. 5 A and D).

After the recording session, small lesions were made at a few recording sites of the silicon probe by current injection, and the silicon probe was drawn from the brain. Cranial window was washed with saline and covered with silicone material. If the animal shows noticeable stress responses (struggling, crying, etc.) to the experimental procedures at any time points, the experiment for the animal was immediately discontinued. The recording session was repeated with the same animal with changing recording sites across several weeks.

### Estimation of recorded sites

After the recordings, animals were deeply anesthetized with urethane and transcardially perfused with saline and 4% paraformaldehyde in phosphate buffer. Exposed brain was coronally sectioned into the 100 μm thickness by vibratome. Recording tracks were verified by fluorescence of DiI under microscopy. Each recording was validated for current source density (CSD) analysis if the recording track penetrates all layers within each area (AON, APC, or PPC) with angle of 90° ± 30° against the layers. Depth position of each channel was estimated using the lesion markings and/or OB, AON or APC stimulation-induced CSD (Ketchum and Haberly, 1993; Takeuchi et al., 2011).

### Data analysis

In this study, we focused on the activity of olfactory cortex during the wakefulness with slow respirations (less than 3 respirations per second). Spike2 software (Cambridge Electronic Design) was used for offline analyses of LFPs and unit activities including power spectrum, coherence, spike sorting, respiration phase histograms and perievent time histograms. To reduce drifting noise of LFPs and respiration traces, DC components of the signals were removed before following analyses.

### Power spectrum and coherence analysis

Power spectrum and coherence analyses were conducted on the respiration traces and the LFPs recorded in the deep layer (200-300 μm deeper to the boundary of the layer I and II) of each olfactory cortex area. We analyzed continuous 100 s data during which the rat showed mostly the wakefulness with slow respirations (over 80% of the time). The power spectrums (from 0 to 100 Hz) and coherences (from 0 to 5 Hz) were calculated at a resolution of 0.1 Hz (200 Hz sampling, 2048 point FFT).

### Detection of slow components of LFPs

For detection of slow activity, LFP recorded in the deep layer of each olfactory cortical area was down sampled to 100 Hz and filtered at 0.5-3 Hz, and the mean value and standard deviation (SD) of the LFP was calculated. When the negative peak amplitude of the slow potential was more than the 2SD, we defined it as depth-negative slow potential (DNSP). If the positive peak amplitude of the slow potential exceeded the 2SD, we defined it as depth-positive slow potentials (DPSP). The timing of the peak of each DNSP or each DPSP was used as the event timing of DNSP or DPSP.

### Current source density analysis

To estimate the positions of synaptic inputs with reference to the layers of the olfactory cortex, CSD was calculated on raw or average depth profiles of LFP recorded at intervals of 100 μm in depth. For the average depth profiles of LFP (Fig. 5 B, C, E and Fig. 7 G), continuous 20-30 s data with very stable respiration pattern were selected and averaged with reference to the onset of exhalations.

The CSD was calculated as a smoothed form from the potentials of consecutive five recording sites with time resolution of 100-200 Hz using a following formula (Freeman and Nicolson, 1975):

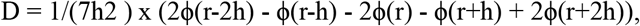

where ϕ(r) was the LFP at depth r and h was the sampling interval (100 μm). The calculated CSDs of each point were linearly interpolated and plotted as a heat map. Hot or cold color represents current sink and source, respectively (Green color is neutral).

### Spike sorting

For spike sorting, recorded data from neighboring 4 sites (50 μm intervals) of linear array was used as a unit of analysis. Single units were isolated using principal components analysis and features of waveforms. We utilized single units which had more than 0.5 Hz of average firing frequency.

### Respiration phase histogram

Inhalation and exhalation phases of respiration were determined by output of the thermocouple which monitored the temperature change in the nostril (Kepecs et al., 2007). Inhalation was detected as the temperature decrease due to the inflow of external cool air into the nostrils. Exhalation was detected as the temperature increase caused by the outflow of internal warmed air. The timing of inhalation onset (the start of temperature decrease) was used to calculate respiration phase histograms. In a respiration phase histogram, one cycle of respiration (from the onset of an inhalation to the onset of the next inhalation) was divided into 36 bins (1 bin = 10°) and the number of events (DNSP, DPSP or spikes) in each bin was calculated. For single unit analysis, the spike activity was expressed as a z-score of each neuron and smoothed using immediate neighboring bins. If the bin of peak activity was in 20-150° or in 230-360°, we classified the data as inhalation or exhalation phase-locked data, respectively.

### Perievent time histogram

To examine the relationship between DNSP/DPSP events and neuronal spike activity, we calculated perievent time histograms (PETHs). Spike activity of each single unit was aligned with the timing of DNSP/DPSP peak (as time zero), and firing frequency (in 0.1 s bin) of the single unit was calculated in the time window from −1 s before the DNSP/DPSP peak to 1 s after the peak. The spike activity was expressed as a z-score of each neuron and smoothed using immediate neighboring bins.

### Statistics

Average data are represented as mean ± SEM. For comparisons of two sample means, two-sided Welch’s t test (Fig.1 D, E, Fig. 5 D-F, Fig. 7 A-C) and paired t test (Fig. 5 D-F, Fig. 6 E) were performed. The uniformity of the phase distribution of DNSPs in respiration phase histograms was tested with Rayleigh test (Fig. 2 I, Fig. 5 G, Fig. 7 D). Sample sizes were justified based on literatures in the field (Sakata and Harris, 2009; Bennett et al., 2013) and no statistical methods were used to predetermine. Data collection and analysis were not randomized nor performed blind to the conditions of the experiments.

## Acknowledgments

We thank members of the Department of Physiology, The University of Tokyo and Laboratory for Systems Molecular Ethology, RIKEN Center for Brain Science for technical advices and helpful discussions. This work was supported by a Grant-in-Aid for Young Scientists (B) (15K16564) to K.N.; a Grants-in-Aid for Scientific Research on Innovative Areas (25135708) and for Young Scientists (B) (25830003) and Takeda Science Foundation to H.M.; a Grant-in-Aid for Scientific Research on Innovative Areas (25115005) to Y.Y.; and a Grant-in-Aid for Scientific Research (A) (23240046) and CREST, JST to K.M.

## Author contributions

K.N. and K.M. designed the experiments, K.N. collected and analyzed the data, and K.N. and K.M. wrote the paper. H.M. and Y.Y. advised the project and critically reviewed the manuscript.

## Competing financial interests

The authors declare no competing financial interests.

## Correspondence

Kensaku Mori

